# A 2-decade Study of Barriers to the Adoption of Organic Farming in Arid Lands of Jordan

**DOI:** 10.64898/2026.02.13.705472

**Authors:** Mohammad Mutarad Al-Oun

## Abstract

Organic farming supports environmental sustainability by saving water, safeguarding ecosystems, and providing economic opportunities through organic crops. It promotes food security and long-term development in the arid regions. However, its adoption in Jordan remains limited, primarily due to insufficient governmental support policies and measures. This study aims to identify the fundamental barriers to the adoption of organic agriculture in Jordan’s arid regions, evaluate farmers’ preparedness for organic practices, explore opportunities for organic farming, and propose recommendations to enhance its adoption. The study utilized a longitudinal approach, conducted in two phases over two decades. The first phase (April–September 2004) involved semi-structured interviews with 46 farmers and five focus groups. The second phase (July–September 2024) revisited seven experienced farmers from the initial cohort, using a phenomenological research approach a widely used approach. The results of phase 1 findings showed that the main barriers were technical, economic, marketing, legislation, institutional and extension and services while socio-cultural was not. The results of phase 2 highlighted persistence of the barriers identified in phase 1, alongside unresolved institutional difficulties, including certification processes, regulatory gaps, and limited market access. The study concluded that implementing streamlined certification procedures, government-supported subsidies, education programs, and policy modifications to promote sustainable adoption of organic farming and farmer engagement in Jorden. The limitation includes a small sample size, the two-decade gap between phases, and a focus on arid regions only. Further, it excludes other stakeholders’ perspectives, underexplores socio-cultural factors and provides limited analysis of certification, market access, and comparable contexts.

## INTRODUCTION

The total area under organic (certified) farming had grown to 96.4 million hectares (ha) by the end of 2022, a notable increase of 26.6% (20.3 million hectares) over 2021, as mentioned in the 25^th^ edition of “The World of organic agriculture” (Willer, Trávníček, & Schlatter, 2024). This increase is due to advanced conventional farming practices and sustainable and eco-friendly farming practices (Panday, Bhusal, Das, & Ghalehgolabbehbahani, 2024). Across the globe, a total of 188 countries practised organic farming and shared 2% of global agricultural land in 2022 (Willer, Schlatter, & Trávníček, 2023). Australia has been reported as the top country with an organic farming area of and 53 million ha, and India has been the second country reaching 4.7 million hectares for organic cultivation (Panday et al., 2024; Willer et al., 2023). Across the globe there are 4.5 million organic producers worldwide, and the estimated cost reached 134.8 billion US$, with significant contributions from the United States and the European Union (Willer et al., 2023).

Along with the global expansion of organic farming, it is essential to under mention the challenges, which include technical, economic, social, cultural, and policy barriers to the broader use of organic practices. In this context, previous reports showed that scalability, certification processes, and profitability are the main problems connected with the development of organic farming (Jouzi et al., 2017; Schneeberger, Darnhofer, & Eder, 2002; Tscharntke, Grass, Wanger, Westphal, & Batáry, 2021). A minimal literature has highlighted the barriers to organic farming in Jorden (Altarawneh, 2016). Therefore, the current study seeks to analyze and provide a basic understanding of the technical, socioeconomic, and cultural aspects of underlying causes for Jordanian farmers. It is, therefore, to provide suggestions for solving the problems in terms of a general or a particular factor. The current study aims to analyze and specify the challenges that hinder the practice of organic farming in the arid zone of Jordan in more detail.

The findings of this study are significant for policy formulation, legislation, research and development, agricultural extension services, market access facilitation, and the encouragement of cost-effective integrated soil and pest management practices in the arid zone of Jordan.

## LITERATURE REVIEW

### Jordan’s Organic Farming and Agricultural Sector

The data published in 2024 showed that Jordan has contributed 1,478 ha for organic farming, as shown in **(Table 1)** (Willer et al., 2022). The reports published by the Food and Agriculture Organization (FAO) showed that Jorden has contributed 264 thousand hectares to total agriculture land (Anbar, Abu-Dalhoum, Maslamani, Al Antary, & Sawwan, 2020) (Agriculture., 2023), as shown in **Table 4**. Therefore, it is estimated that Jorden has a very limited contribution of up to ≤1 % of organic farming in total agricultural land (Willer et al., 2022). More interestingly, the data shows an increase of 32 ha between 2021 and 2022. However, there has been a decline of 1,420 ha in the last decade. Unfortunately, not a single piece of literature has discussed the reasons for the decline; therefore, the current study will highlight a few reasons based on the results.

In this context, **Table 2** shows Jordan’s socioeconomic key organic indicators. These indicators show that Jordan has only 16 producers and seven processors. The country exported 86 metric tons of organic items, 70 metric tons of which were exported to the EU, and 16 metric tons of coffee to the USA, even though Jordan is not a coffee-producing country. Although the organic farming area is small, Jordan has a diver’s organic crop production: cereals, temperate fruit, Tropical/Subtropical fruit, Grapes, Olives, and Vegetables.

**Figure 1** shows the inconsistencies in Jordanian land for organic farming between 2009 and 2022. The area was 3,500 ha in 2009 but declined to 1,478 ha in 2022. A more significant decline is observable from 2013 and onwards, with a plateau from 2016 to approximately 1,400 ha where an increase of 32 ha in 2022 shows some consistent difficulties in maintaining organic culture.

**Figure 1:**
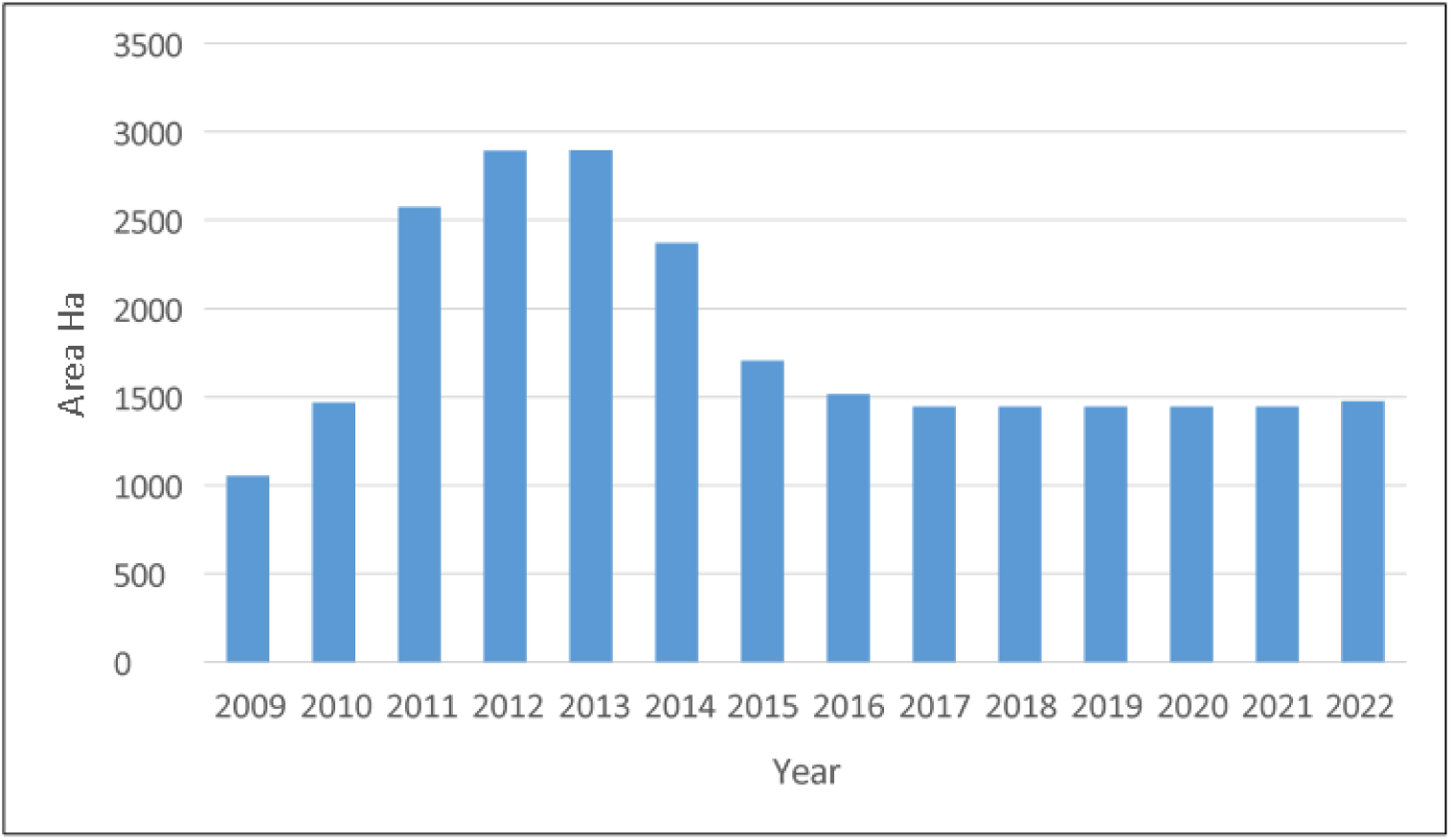
Change in Jordan’s organic farming land over 14 years (Data derived from FiBL and IFOAM reports 2014-2024).

A comparison of some key organic indicators in Jordan **(Table 3)** with those of neighbouring countries, such as Saudi Arabia. Saudi Arabia’s organic farming area is 23,315 ha, more than 10 times larger than Jordan’s. Moreover, Saudi Arabia has 512 organic producers, whereas Jordan has only 16. A comparison with the United Arab Emirates (UAE) shows that the UAE has 5,419 hectares and 152 producers, exceeding Jordan’s figures. Despite this, Jordan’s conventional data shows that the country has diverse crops, which would allow it to convert some conventional agricultural areas into organic areas, as shown in **Table 4**.

### Diverse Barriers to Adoption

The barriers inhibiting the widespread adoption of organic farming encompass an extensive spectrum of challenging factors. These obstacles include but are not limited to restricted market access, inadequate basic experience, high costs of acquiring organic certification, and high capital investment requirements (Dixit, Suvadarshini, & Pagare, 2024). The frequent barriers include socio-cultural and legal structures, which are crucial factors in embracing change. The challenges include labor issues that differ substantially between developed nations and the developing world. Therefore, innovative strategies are highly required to cope with the challenges in different conditions (Łuczka & Kalinowski, 2020; Sharifi et al., 2010). In addition, factors that affect literacy levels, such as low literacy levels, also hinder the farmer’s activities and complicate the situation (Zhang & Wang, 2024)

### Regional Specificities in Barriers

The trends, patterns, and intensities of barriers are dissimilar across regions and thus require place-based approaches. For instance, capital costs, which include expenses on certification, are significant challenges to smallholders planning to do organic exports in different areas of Africa, or the high costs of accreditation and lack of sufficient rules that hamper organic farming in South Africa (Eyinade & Akharume, 2018; Mkhize & Ellis, 2020). Moreover, scarce infrastructures, fluctuation in the market indicators, and lack of proper advisory services also aggravate the problems, especially in some regions (Oghazi & Mostaghel, 2018).

### The Imperative of Investigating Barriers

It is inevitable to understand the farming system; therefore, organic farming requires commitment and hard work before suggesting and adopting an organic farming system (Eyhorn, Van den Berg, Decock, Maat, & Srivastava, 2018; Reganold & Wachter, 2016; Schneeberger et al., 2002). In case of failure, these barriers can hamper finance and operation issues during the conversion process, which may seriously threaten organic farming initiatives regarding sustainability and success(Eyhorn et al., 2018; Łuczka & Kalinowski, 2020).

### Organic Farming in Jordan: Contextual Challenges

Jordan being a developing country, realizes that it has major challenges in the agriculture industry due to climatic and economic factors of the country. The country has an economy of $49.29 billion and $4,311 per capita; unemployment is expected to be 21.4% in 2021 (Bank, 2022; Economics, 2021; News, 2022). Water scarcity is very serious in Jordan and this has a very negative impact on agriculture that is already a candidate for climate change because of the climatic change from Mediterranean type to desert type. Due to scarce natural resource base, high dependence on traditional irrigation techniques and water dependent cropping systems, maintaining the productivity is a function of synergistic and flexible practices (de Vito, Portoghese, Pagano, Fratino, & Vurro, 2017). Also, since Jordan critically relies on imported food to feed the citizens, this increases the concern for the country’s insecure agricultural sector and the necessity to invest in resource-saving approaches.

Organic farming holds out the potential to provide such a solution to these issues: sustaining action and conservation of resources. But certification is still low due to demerits such as high costs in certification as well as low marketing for organic food (Fuhrmann-Aoyagi, Miura, & Watanabe, 2024). Organic farming practices can be accepted within the existing model of extensive agriculture and policies promoting such strategies can encourage producers to work more closely with the natural environment in Jordan and to develop a locally resilient, sustainable food supply system. Achieving these solutions will therefore require a policy and investment approach to eliminate the barriers that have been observed as inhibiting development of organic agriculture into the foundation of the nation’s agriculture.

## MATERIALS AND METHODS

### Study Area

The current study was carried out in the North-East Badia (NEB) of the Mafraq Governorate, a region in Jordan that the Soil Survey of Mafraq Governorate categorized as having organic agriculture climate ((MoA 1994). On average, this area has temperatures of 17.5°C per year, with minimum temperatures that range between 10 °C during the winter season and maximum values that exceed 46 °C in the summer {Allison, 1998 #98}(Allison *et al*. 1998). The annual rainfall is estimated to be around 200mm, and as a result, adoption of sophisticated methods of irrigation, such as drip irrigation. The area mainly adopted cash crops agriculture; the climate and farming practices offer a realistic environment to related difficulties and potentials of organic (Portal, 2021) as shown on the map in Figure 2, maps and locate the geographical location and context of the study area.

**Figure 2:**
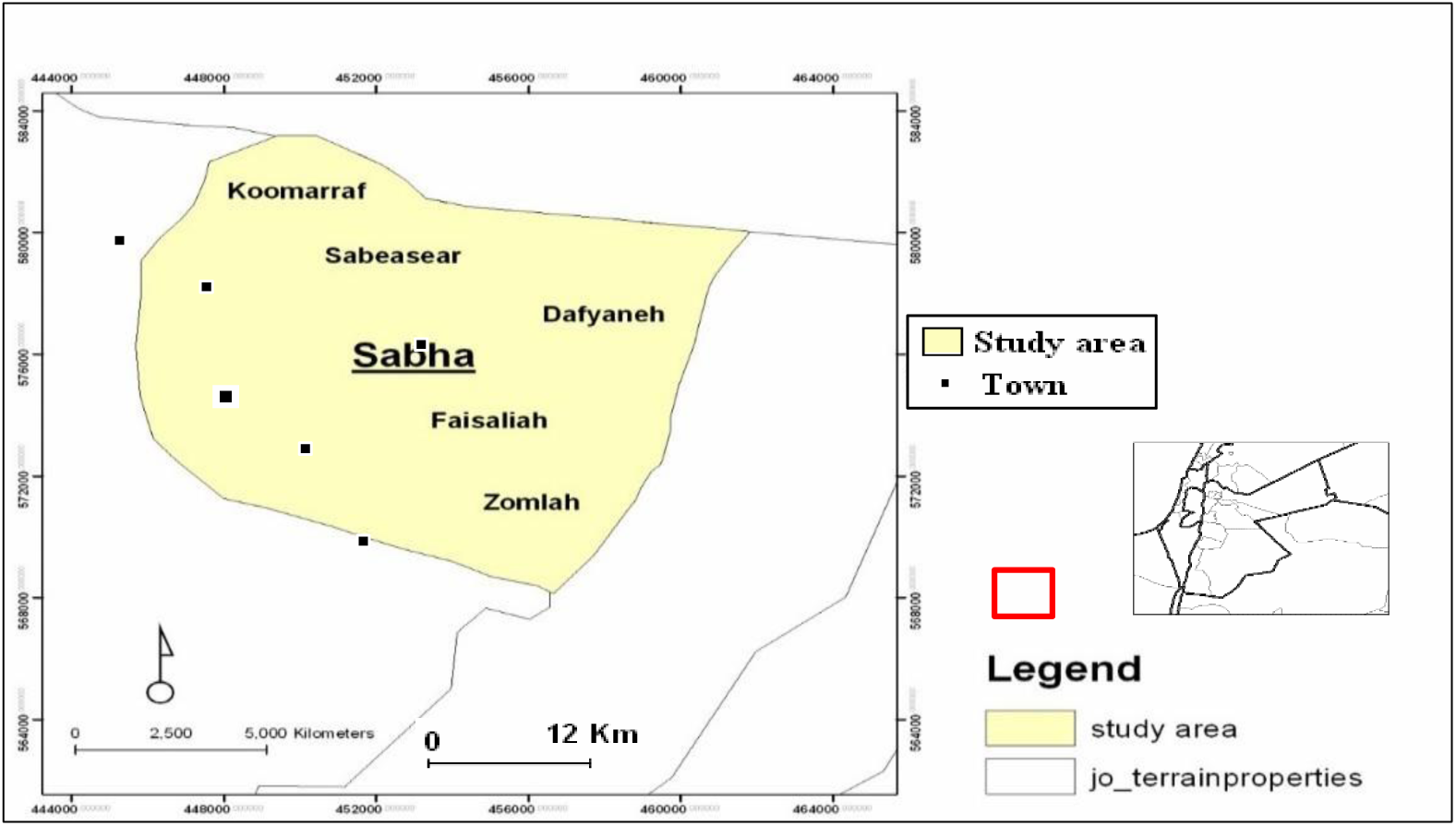
The location of the study area (Modified from JBRDC, 1994).

### Research Design and Approach

The research used qualitative and quantitative data collection methods to capture all the factors that may hinder the adoption of the organic farming system. This approach was implemented in two phases over two decades:

- Phase One (April and September 2004): Primary data collection was done using the exploratory research design, 46 farmers and five focus groups were interviewed using semi-structured interviews. Also farms information (types and coordinates) were collected to produce a map (Figure 3) showing their types and distribution.
- Phase Two (July and September 2004): Subsequent studies will use the same set of farmers (where reachable) to gauge the evolution of barriers and perceptions towards getting into and maintaining a fully organic farming practice.

**Figure 3:**
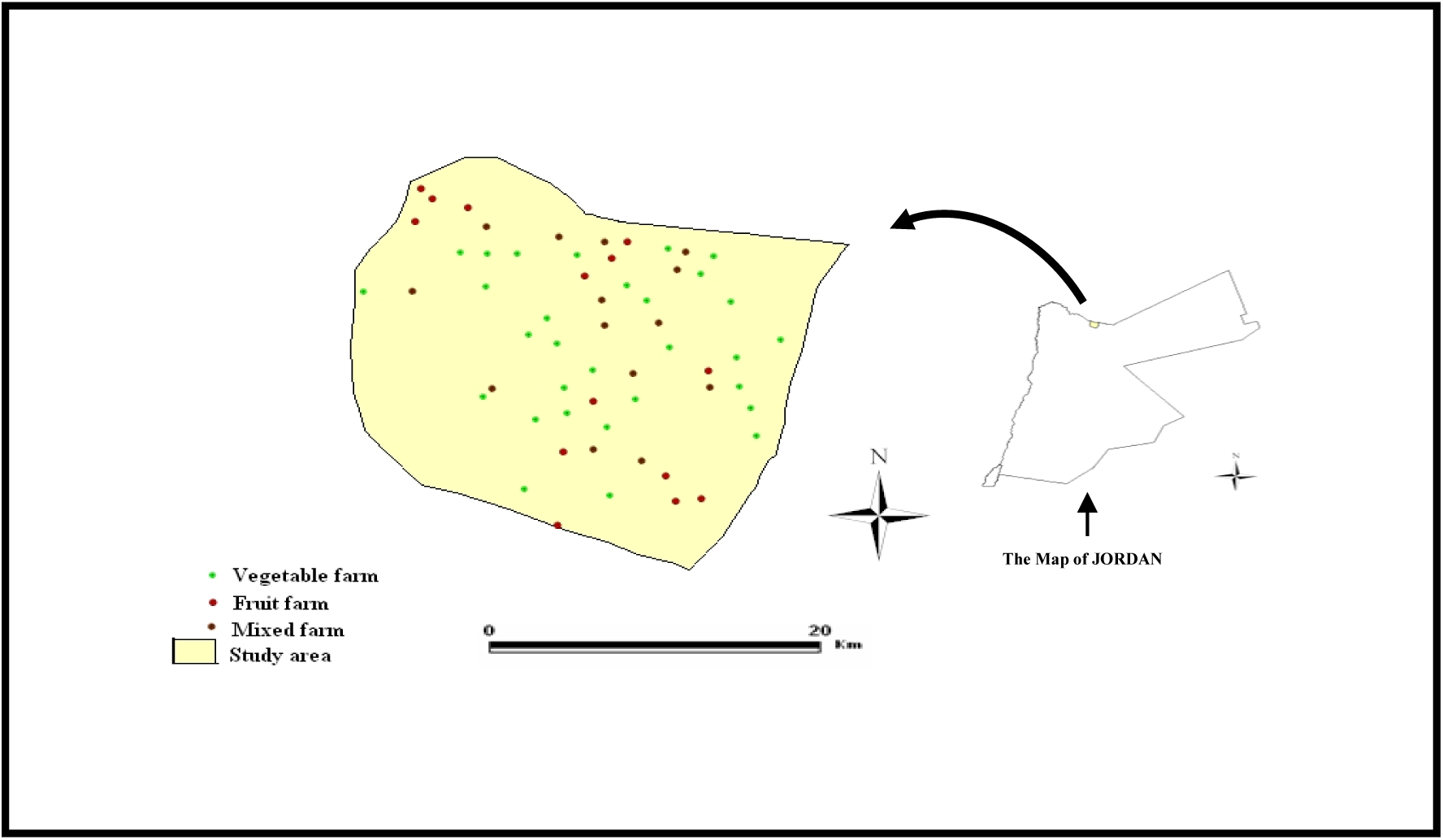
The study area farms’ types.

### Data collection strategy

The approach used to collect the primary data in Phase One used in this research was informed by two empirical research works (Crowder & Illan, 2021; Schneeberger et al., 2002). The first study looked at the hurdles constraining organic farming adoption among Austrian cash crop farmers and adopted a mailed survey questionnaire on 1,000 farms, of which 383 responded. These responses identified 28 potential challenges and solutions for increasing numbers of organic producers. The second study targeted the SSA region using a postal survey of 213 community development groups across 24 countries and field research in Ghana and Kenya. Focal group discussions and interviews further complement the information gathered from three agro-ecological zones.

The current study followed this approach (Crowder & Illan, 2021; Crowder, Northfield, Strand, & Snyder, 2010; Krause et al., 2024; Muller et al., 2017) to investigate barriers in detail in the Jordanian context and utilised multiple data collection techniques for enhanced understanding of the barriers.

### A postal survey or semi-structured interviews

In Jordan, physical data collection methods, particularly postal ones, are not feasible to implement because of barriers, including the inefficiency of mail deliveries, faxes, and fixed addresses. To overcome these deficits, this research used a semi-structured interview method suggested by other scholars for similar settings (Crowder & Illan, 2021; Schneeberger et al., 2002). Semi-structured interviews proved more appropriate to cover some sensitive issues, such as cropping systems and the opportunities and constraints of organic farming, than postal surveys, which are likely to have low response rates and incomplete data (Crowder et al., 2010).

### Bottom-Up Approach and Data Collection

This study used the bottom-up approach, focusing on the farmer as a user, major player, and end consumer in the agricultural value chain. According to (Muller et al., 2017), involving farmers right from formulating research questions ensures that the results and recommendations provide solutions to the ground needs rather than what researchers or policymakers perceive. However, this approach also has drawbacks as part of the relevance and inclusiveness sought to be achieved. For instance, in organizations with no institutional support for participatory systems, integration can be a challenge (Fantinelli, Di Fiore, Marzuoli, & Galanti, 2022).

Furthermore, adopting this farmer–acquired information may not capture other general and environmental factors affecting farming (Crowder et al., 2010). This is why it is necessary to combine participation findings with scientific evidence to prevent the emergence of too-district or stringent outcomes. Implementing the participatory methods is also hampered by the limitations in funding and other resources. Therefore, enhanced institutional frameworks and harmonized agricultural policies are essential.

However, with proper backing of institutional frameworks and coordination with general agriculture policies, a bottom-up approach can be a strong instrument for collecting valuable data.

### Data Coding and Entry

Following the completion of fieldwork, responses were categorized based on questions in a codebook, assigning numeric codes to each category for the systematic organization (Carey, Morgan, & Oxtoby, 1996; Rose & Sullivan, 1993). Subsequently, these coded responses were manually inputted into the Statistical Package for Windows (SPSS) to create variables and ensure data entry accuracy (Carey et al., 1996; Rose & Sullivan, 1993; Walz, 1999).

### Descriptive Analysis

Qualitative data was analyzed using descriptive-analytical methods to code, categorize, and summarize the results. The findings were summarized in tables using frequency analysis, which gave a clear output of the perception of the interviewees and the constraints on organic farming.

### Phase Two: Assessment of the barriers after 20 years

Between July and September 2024, semi-structured interviews were conducted with representative farmers (participants) to establish whether there were changes in the barriers to implementing organic farming in the study area, whether the obstacles remained the same or if new barriers had cropped up. The phenomenological research approach was used because this approach is appropriate for exploring participants’ experiences (Crowder & Illan, 2021; Tarawneh, 2021). Although this approach was used to reduce generalization, it was effective in understanding the local farmers’ experiences.

The study employed purposeful sampling to select seven participants with vast experience in farming in the study region. These participants offered the study a wealth of understanding of what goes on in agricultural practices and the problems they face. These barriers described in Phase One include perception, technical, cultural and social, marketing and economic, institutional, regulatory and legislative, information, advisory, and services barriers for the follow-up investigation. Since there hasn’t been any literature explaining the reasons behind the decline, as shown in Figure 1, therefore, the farmers were invited to share their views on the matter. Semi-structured interviews were arranged to gather more detailed contextual information and avoid artificial and narrow-folded answers.

### Thematic Analysis

Data was analyzed thematically to pinpoint nuanced patterns and trends surrounding organic farming adoption based on the interviews conducted. This included using simple templates to transcribe interviews, categorising interview answers into themes and synthesising themes to identify key issues. The identified issues included a lack of proper control mechanisms for pests, few organic certifications for crop production, low soil fertility management, and fewer market outlets for organic crops.

However, because the study adopted a case study approach, the sample size was comparatively small, encompassing only seven farmers. Generalizing the results to other farming regions was thus a challenge. The effect of different types of crops grown for varied durations was not captured precisely: Short-duration crops grown by the vegetable farmers and long-duration crops grown by the fruit tree farmers. Other factors such as mixed farming activities, farm size, and farming experience were other causes of variations that could have been mitigated but not well handled.

Although the purposive sampling approach effectively provided depth in the study of the researched phenomenon (Tarawneh, 2021; Tscharntke et al., 2021), the limits of this study should raise questions for subsequent research with more diverse populations. A larger variability of farms regarding size, gender, age or region using a stratified sampling method could give further insight into the problem. By engaging stakeholders such as policymakers, agricultural scientists, extension agents, retailers, and consumers, the study would adopt a much wider lens to view structural, institutional, and market impediments to organic farming in Jordan (Muller et al., 2017).

### Research Ethics and Confidentiality

Regional ethics shall be strictly ensured during both study phases. Potential participants were told about the purpose and nature of the study and then gave their consent before involvement. They were told they had the right to refuse to continue with the study without explanation, and their answers would always remain private. (Rose & Sullivan, 1993). Consent forms declared that results would be published without using the participants’ identifiable information, including their names.

The study did not require data from participants, nor did it use personal information and/or direct interventional procedures; therefore, it did not call for the approval of an ethics committee. In this way, it was possible to safeguard the general principles of ethical research and the anonymity of participants simultaneously.

### Strengths and Limitations

One of the significant strengths of this study is that the research activity focused on farmers across two phases: five groups and seven interviews, which helped identify key challenges for adopting organic farming. It examined various issues and provided commentary regarding the prospects of organic farming production in the area.

Nevertheless, some of the study’s shortcomings include a few survey participants and geographical constraints that limit generalizability. Subsequent research should try to include specific solutions in addition to the accounts and assess the thenal impact of those solutions in the long run.

## RESULTS AND DISCUSSION (Phase One)

### Farm types and farmers socio-demographic characteristics

Results showed that there were 57 farms, comprised of 14 fruit farms, 14 mixed farms (vegetables and fruit), and 29 vegetable farms, as shown in Figure 3, with a total land area of 2153 ha, as shown in Table 8, but only 1486.6 ha were actively cultivated. The average farm size was 37.1 ha, while the average cultivated area per farm was 25.7 ha, indicating that approximately 30% of the land remained uncultivated for crop rotation purposes. Vegetable farms occupied the largest proportion of the cultivated area (37%), followed by mixed farms (36%) and fruit farms (27%).

The research analyzed the socio-demographic characteristics of 46 male farmers (average age of 51 years) in the study area, showing a diverse range of educational backgrounds, as shown in Table 9. Furthermore, farmers who owned a single farm 79%, while 24% were not originally farming in the study area, indicating a shift in the local community structure. This study observed a strong social bond in the farmers’ community, therefore, considering it is potential for collective action, such as in terms of adopting organic farming practices.

### Farmers’ perceptions

In the case of organic farming, it was identified that only 35% of the farmers participating in the sample had ever heard about or knew some information about organic farming. The activities depended on and aligned into two categories. They were divided into two groups.

- First, 28% had heard about organic farming by attending three workshops. This group of farmers only knew that organic farming was about growing crops without applying pesticides and fertilizers (leaving chemicals) and using cow dung and water. They realized from the workshops that to be ‘organic,’ some regulations must be complied with, but apart from that, they were hardly aware of many things about organic farming.
- Second: 7% of three farmers had heard or knew of organic farming through their work. Those farmers were a University Vice Chancellor, Ministry of Agriculture Ex-Secretary General, and an Agricultural Engineer fruit exporter to Europe.

On the other hand, 65% responded that they had never heard about organic farming but provided different expectations and opinions, as follows:

- Organic farming means the use of manure

This perception was reported by 35% of the farmers. It was noted that these farmers linked the words ‘organic’ and ‘manure.’ The translation for organic to Arabic is alodweyah; also, manure means, in Arabic, ‘organic fertilizers’ (Alasmedah alodweyah). This explains why farmers thought that organic farming meant using manure. This also was the perception of 60% of the focus groups who were well-educated in agriculture.

- I have no idea about organic farming

In 3% of the farmers, reported that their first hearing about organic farming from this research.

- Organic farming means the minimum use of chemicals, such as integrated pest management products.

6.5% of the farmers reported that organic farming means the minimum use of chemicals, which means the IPM.

- Organic farming means Baeel

11% of the farmers reported this. Baeel is an old traditional agricultural system in which farmers used to depend on rainfall or any water source to water their crops and use manure as a fertilizer. In this system, no chemicals were used except sulfur. It is a non-certified organic farming system.

### Technical Barriers to Organic Farming-

The study aimed to discern potential technical barriers farmers anticipate regarding adopting organic farming practices in Jordan’s arid lands. The reported barriers were cross-referenced with existing literature, revealing ten notable technical barriers (Dixit, Suvadarshini, & Pagare, 2022; Läpple & Kelley, 2013), as outlined in **Table 5**. These highlighted barriers echo prior studies while unveiling context-specific factors unique to this research.

Consequently, **Table 5** shows the technical challenges to organic farming as captured by 46 farmers and five focus group discussion sessions. The most reported difficulties include higher disease and pest incidence and lower yield quantity and quality, each of which was cited by all respondents. Low crop yield, stunted plant growth, and poor soil quality were other problems mentioned by more than 90% of the sample. Other challenges include the danger of using the organic method (70%), appropriate plant varieties (61%), and access to biocontrol agents (61%). Poor periods of production (57%) and more focus on their expertise instead of farmers’ (50%) were seen as a problem as well as changing weather conditions (28%).

### Cultural and social

The Jordanian farmers are not restricted by culture or social norms in the practice of organic farming or conventional agriculture since their main aim and objective is to realize financial gains from the production of crops. Interestingly, the intergenerational conflict and the fear of social isolation that a study mentioned were not seen as obstacles by respondents (farmers)(Padel & Lampkin, 2007). Consistent with Krause *et al.,* this study found that the fear of failure was a significant concern raised by the farmers, who feared social backlash in case of failure in farming (Krause et al., 2024). This fear, due to perceived social status and the desire to be recognized as a ‘successful farmer,’ discouraged using the technology, which was considered a risky and unproven farming method. Although the cultures of the two groups did not forbid the production of organic produce, the need to sustain a noble image in the eyes of their people was established to be a factor that discouraged organic farming. In their assessment, Krause et al., found that factors that farmers in other regions fear of social censure of failure also affect their farming. This explains why they avoid adopting other practices, like organic farming, which they regard as unproven new-age fashion (Krause et al., 2024). While farmers mentioned that consumers may have cultural taboos against consuming organic products, farmers disagreed that they also had cultural taboos against producing them. One of the study’s main findings is that social fear of failure is one of the key factors that prevent the adoption arising from individuals’ desire to maintain a good reputation among their peers.

### Economic Barriers

Farmers expressed concerns about the need for a market structure conducive to organic products and the financial strain associated with transitioning to organic practices. The need for premium pricing avenues for organic goods significantly hindered their ability to offset increased production costs. This financial strain and substantial farm investments raised doubts about the profitability of organic farming endeavors, as mentioned in **Table 6**. These economic hurdles significantly deter farmers’ perceptions of the viability and sustainability of organic farming practices. **Table 6** presents different frequencies that show the effect of economic barriers on the implementation of organic farming, with percentages ranging from 52.2% to 97.8%. The lack of a market or high costs are also linked to the highest frequency, indicating a serious issue. On the other hand, financial commitments and possible unemployment show much lower frequencies, indicating fewer challenges in these domains.

### Marketing

In this study, five major marketing problems affecting organic farming in Jordan are highlighted. As per the study, 95% of respondents reported that customers and consumers lack of understanding of organic food as a challenge. This was succeeded by a lack of efficient organic marketing communication networks (90%), inadequate premium pricing (90%), and geographical space between the producers on the one hand and the markets or delivery points on the other hand (80%). Farmers are convinced that there is not enough demand for organic produce and that conventional products will compromise the sales of organic produce. This concern is similar to what studies have found (Lampkin, Padel, & Foster, 2000). For example, the farmers mentioned that previously there was a company specialized in marketing integrated pest management (IPM) vegetables products who received much support from the government and international organisations but failed to popularize of its vegetable products, due to the market’s lack of appreciation for those products; the company sold over 98% of them as conventional products. The company could not sell more than 1,000 kg of IPM products daily, which was never enough to bring at least one employee to the shop. Consequently, the IPM company barely made profits and dismally only met its expenses, forcing it to switch to conventional products. When implementing the mentioned IPM project, the company learned that this production system had no market. Accordingly, selling organic products is a high-risk business; therefore, they-had no plan to encourage organic farming. One farmer said, “If the organic products are introduced to the market, they will end up being like the IPM products and sold as conventional products. All respondents noted that the lack of understanding of the organic farming processes is a barrier to firms adopting those processes because the consumers are only familiar with the end product. This postscript supports Walz’s (1999) assertion that marketing is hampered by low consumer knowledge of organic food and underdeveloped marketing channels. Also, the findings support the findings of Schneeberger, Darnhofer, and Eder (2002), who pointed out that the absence of marketing channels coupled with difficulties in obtaining high prices for the products are common impediments to the organic farmers’ conventional counterparts.

### Institutional

For a detailed analysis of the institutional factors affecting the growth of organic farming, refer to (Al-Bitar, 2006; Padel & Lampkin, 2007). Padel and Lampkin et al. (2007), have identified some barriers to organic farming as being the refusal of landowners to switch to organic agriculture, financial institutions’ inability to support or underwrite crop farming organically, difficulties in accessing grants, legislation that inhibits organic farming, and problems with certification (Padel & Lampkin, 2007). Whilst this study has established some barriers, others were not emphasized; this shows that there is a need to develop more institutions to function in the agricultural sector within the Jordanian context, including the Ministry of Agriculture (MoA), National Agricultural Research Center (NARC), Agricultural Credit Corporation (ACC), and universities. However, these existing organizations should enhance their cooperation. They embrace legal services, extension services, research, credit, and agricultural services to the farming community.

Therefore, the first significant challenge is that none of these institutions have long-term policies that endorse organic farming. Specific policy support for organic agriculture is still missing, which is one of the critical issues Al-Bitar (2006) mentioned for the Arab region. Out of all the interactions made with individuals from such institutions, another issue is the failure of institutions and their target groups to synchronize themselves and, consequently, failure to address key proposals. For instance, the study revealed that two institutions may perform or fund activities in the same thematic fields without coordination. Without consulting, these institutions quickly make new policies or present proposals that affect farmers. For example, an observed conflict of interest emerged between the staff of the organic farming unit (OFU) and the mandate of the MoA. Conversation with the Secretary General of the MoA and the head of the MoA Policy Directorate found that the Jordan Institution for Standards and Metrology (JISM), with the support of other ministries responsible for setting the standard of all different food products of Jordan, including the organic farming business. However, the OFU has also claimed that the MoA can set these standards.

Two examples are the exclusion of the NARC from the analyzed methods in organic farming and the fact that OFU did not seek collaboration with the JISM. Also, producers were not included in the MoA’s plans to promote organic agriculture for food producers. The MoA Secretary-General had stated that there were plans for developing organic olive oil and herbal plants, but they found that farmers had never been consulted. The OFU was concerned only with olives. That is why the urge for organic farming necessitates coordination between these institutions, not replication. This must be easy due to the sound and effective communication that the above organizations engross, and there should be an overarching body to coordinate them.

Another area to do with credit facilities, as much as the primary concern of high-interest rates, is that organic farmers are declined loans. At equal measure, there are no insurance policies for organic farming. It demonstrates that, in general, loans are offered for conventional systems of production rather than for organic methods. Hear for it now from the farmers again: while the ACC used to fund traditional agriculture, for instance, US$60000 for good digging, water harvesting projects, which cost US$7000, are no longer entertained. The fourth challenge is that organic farming is well understood more as a production technique than a complete production system. Thus, the difficulty in talking about its use, let alone viewing it as a separate anything in MoA, is because, as rightly pointed out by the MoA Policy Director, organic farming is another method within crop husbandry.

### National regulation and legislation

Specific rules are mandatory prerequisites to organic farming’s existence. Regulation helps to provide the transparency necessary for the smooth functioning of agriculture. This framework is essential for consumer confidence and market development (Willer & Yussefi, 2000). According to Walz et al., the greatest obstacle to organic farming is the lack of national legislation (Walz, 1999).

Likewise, product certification and the organic farming and production industry face a significant challenge with poor regulation and legislation in Jordan. In this research exploring the newly developing sector, I realized that no legal protection is given to supporting organic agriculture. Though Jordan has Agricultural Law No. 44 of 2002, which deals with all aspects of agriculture: plant production, fertilizers, pesticides, livestock health, fisheries, etc., Jordan has no specific regulation regarding organic farming or the use and labeling of organic produce. The law says in Article 1 that it shall be called The Law of Agriculture for 2002 and come into force on the 30th day from its publication in the official gazette. The law has 73 articles, but there are no law provisions regarding the methods used to produce the products or the labeling of the products. In the market for agriculture products, the unique aspect is quality and where it is from, such as the southern and northern regions of Jordan or the Jordan Valley.

They also noted that the case of Jordan shows that this country has no particular regulations on organic farming, or at least on the certification and inspection of products. When asked, all the farmers participating in the research and the MoA officials boldly noted that lack of regulation hinders the growth and development of the organic farming sector. Similar to the current attitudes expressed above, the Jordan food standards officer from the Jordan Institution for Standards and Metrology (JISM) and a University Vice Chancellor announced that, at present, Jordan lacks a system for regulating organic farming starting with the laws observing the international standards. All respondents across the study identified that the main challenges to the practice and adoption of organic agriculture are the absence of well-developed national codes and certification programs. This issue is also evidenced in the literature on organic farming in the Arab world, which points out that this is a reason for the lack of regulations (Al-Bitar, 2006; Walz, 1999).

### Information, advice, and services

This literature review reveals the fact that the absence of adequate information and directions was one of the factors that hindered the application of organic farming in Jordan. This finding supports the previous studies (Darnhofer & Schneeberger, 2007; Padel & Lampkin, 2007; Schneeberger et al., 2002). Since organic farming has very low existent in Jordan, the duty of providing advisory and extension services, as well as the national media and official bodies of agriculture, is limited to conventional farming practices. Likewise, the present study for organic farming found that 98% of farmers and the focus group participants had no access to organic farming information or extension services. Some of the points made by the participants include the fact that it takes farmers and extension agents equal time to develop an understanding of organic farming. They also agreed that while the absence of information is an enormous problem, the lack of knowledge among the extension agents is equally problematic.

Besides, it was noted that all the extension materials and advisory MoA leaflets are made using traditional farming techniques. Furthermore, a University Vice Chancellor complained that, according to his knowledge, no university in Jordan is researching organic farming as an integrated system. Most research has concentrated on a particular soil type or pest problem, often in isolation. However, nobody has taken a systems approach to looking at the workings of organic farming.

### Potential Opportunities in Organic Product Production

Both farmers and the five focus groups (private agricultural store suppliers) highlighted the study area’s potential for producing organic products, as presented in **Table 7**. Recognizing untapped opportunities aligns with broader sentiments regarding the region’s suitability for cultivating organic goods. This acknowledgement of potential signifies an area ripe for exploration and development within organic farming. **Table 7** displays the frequencies of the focused group; this table gives insights into the alignment of focused groups with categories including Wide area and available virgin lands,” “Availability of water with good quality,” and “Long production season,” which together account for 25% of all the groups. Furthermore, 20% of the focus groups are categorized into categories like “Good weather and not polluted” and “Soil is not polluted,” but 10% are assigned to categories like “Livestock present,” “Labour (workers) available,” and “The biggest tomato factory in the country.” Finally, 15% of the focus groups included the statement, “Farmers accept new profitable ideas.” The economic, institutional, and regulatory challenges surrounding implementing organic farming practices in Jordan pose multifaceted hurdles. Strategic interventions, collaborative engagements, and establishing standardized protocols are imperative to bridge existing gaps. A cohesive ecosystem involving regulatory bodies, research institutions, and agricultural stakeholders is crucial to unlocking latent potential and fostering sustainable growth in the organic agriculture sector in Jordan.

### Respondents’ Rating of Barriers to Organic Farming

Without existing literature on certified organic farming practices specific to Jordan, this study compared its results with global and local insights (Gamage et al., 2023; Rose & Sullivan, 1993; Smith et al., 2019; Trukhachev, Belopukhov, Grigoryeva, & Dmitrevskaya, 2024). Six categories were created from the interviewees’ challenges: perception, technological, financial, cultural, promotion, and institutional. Notably, barriers related to animal production were not extensively discussed, although the research highlighted livestock as a significant barrier, while labor was not considered problematic. Surprisingly, none of the farmers identified labor as a potential factor, indicating that acquiring local or external laborers was not a significant challenge. While previous studies have explored technical barriers to adopting organic farming, this research uniquely investigates these barriers based on conventional farmers’ expectations.

### Perception as a Significant Barrier

Perception emerged as a significant barrier, resonating with previous research findings (Alotaibi, Yoder, Brennan, & Kassem, 2021; Da Silveira, Da Silva, Machado, Barbedo, & Amaral, 2023). Surprisingly, the term ‘organic farming’ needed a more precise definition among respondents, including Ministry of Agriculture officials. This ambiguity led to varied interpretations among farmers. For example, while a small proportion associated organic farming with Integrated Pest Management (IPM), only 30% associated it mainly with manure. To be more specific, 20% saw it as a “Baeel” production system that entirely depended on rain and did not embrace the application of any chemicals except for sulfur.

These diverse perceptions were due to the lack of prior studies concerning the term ‘organic farming’ in Jordan. The farmers easily related organic farming to manure because the Arabic word for organic resembled the word for manures. Thus, this correctly understood notion prevailed among the focused participants in agriculture, but among the educated people, it was erroneous.

Furthermore, some farmers equated organic farming with the least chemical input, as in the case of Integrated Pest Management (IPM). The previously dominant understanding of ‘Baeel’ and rainfed, low-input agriculture using chemicals sparingly and sulfur has become the most widely held perception of non-certified organic farming systems.

### Implications for Organic Farming Adoption

The results confirm the importance and the necessity of establishing a precise and unified definition of ‘organic farming’ for the farmer and other related parties. The misconceptions and various interpretations showed that providing a clear and coherent understanding of organic farming is necessary. Dispelling these misconceptions and raising awareness is critical to creating the conditions necessary to encourage the successful adoption of organic farming in Jordan.

### Impact of Pest Infestation and Yield Reductions

A significant drawback of pest outbreaks and a probable decline in yields is another challenge farmers experience when switching to the organic farming sector. These fears are not unfounded since Schneeberger et al. note that pests, insects, diseases, and weeds have become more challenging to control compared to those without synthetic pesticides (Schneeberger et al., 2002). The latter is often characteristic of organic farming systems, which by definition do not use conventional chemical treatments and, therefore, aggravate these problems. Crowder and Illan et al., opined that the change to organic practices, especially when using compost, animal manure, and crop rotations, may increase pest pressures, resulting in the expected declines in yield size and quality (Crowder & Illan, 2021). Pesticides were identified as amenity crops, and farmers were concerned that lack of pesticides would aggravate pest menace, which would be detrimental to maximum yield. This leads to the situation whereby organic methods, as sufficient as they may be, hamper production and cannot be implemented on a large scale without complementary pest eradication and enhanced farming techniques.

### Impacts on Plant Growth and Soil Fertility

Expect a decline in plant growth due to lack of nutrients and continuous pest attacks as a significant challenge. Close to all farmers exhibited concerns about several factors affecting plant growth, a key factor in production among farming communities. Likewise, the issues with low soil productivity in the Badia region have prompted the attention of the farmers, where 91 percent considered it a significant technical challenge. The acute necessity to use fertilizers to maintain viable crop production was vividly pointed out, as well as the capabilities of the soil to provide the necessary nutrients for plant growth and development.

### Risks Associated with Organic Farming

Some of the farmers’ concerns concerned the inherent risks of using organic farming. About 70% of the respondents perceived the venture into organic farming in the Badia region as highly risky. This is due to a reluctance to take risks and proactive chemical spraying despite no sighting of pests seen as optimal. The resistance to abandoning chemical fertilizers and pest control and adopting organic farming is due to the risk associated with the unknown and yield reduction with organic agriculture.

### Challenges with Crop Varieties and Pest Control Options

The fact that most conventional crop varieties are unsuitable for organics due to their high vulnerability to pests became identified as a significant challenge, in line with issues raised by 61% of farmers. The lack of effective competitors to pest control, especially the impossibility of biological control in open-field farming in Jordan, became challenging. Private stores and government agricultural departments’ examinations of organic material were scarce. This scarcity gave the farmers more reasons to fear embracing organic farming practices.

### Impact on Production Timelines and Knowledge Acquisition

A substantial proportion (57%) of farmers foresaw prolonged production timelines as an inevitable consequence of embracing organic farming. Ceasing the usage of fertilizers and insecticides as integral components of organic farming was thought as a catalyst for extended production cycles; this transition, promising environmentally friendly outcomes, was projected to yield lower quality and reduced crop quantities over prolonged periods, triggering potential financial losses (Holka, Kowalska, & Jakubowska, 2022). Additionally, concerns were raised regarding abandoning conventional farming experiences, leading to a steep learning curve and uncertain outcomes.

### Weather Fluctuations as a Minor Barrier

Despite the relatively minor emphasis, some farmers (28%) acknowledged variations in weather specifically as a possible technical issue during the end of April and the beginning of May. While only two focus groups deemed weather as significantly challenging to organic farming, this aspect merits consideration due to its intermittent impact on crop production.

### Institutional Barriers

The fragmented landscape of stakeholders involved in organic farming unveiled institutional challenges. The lack of cooperation among key organizations, such as the National Center for Agricultural Research (NARC) and the Jordan Institution for Standards and Metrology (JISM), presents a systemic challenge. Additionally, the Ministry of Agriculture’s (MoA) drive towards organic farming has excluded farmers from participating, underscored by a disconnect among stakeholders, hindering concerted efforts to promote organic practices.

### Regulatory Challenges

Jordan’s absence of established national organic standards or a comprehensive inspection and certification system for organic farming practices was identified as a regulatory challenge. This regulatory gap complicates the landscape, emphasizing the need for structured guidelines and oversight mechanisms to effectively control and promote organic farming practices.

## RESULTS AND DISCUSSION

### Phase Two

Phase two of the study was to determine whether obstacles to implementing organic farming, established in Phase one twenty years earlier, are still considered a problem by farmers. In detail, it looked at sustenance issues and investigated how these hurdles influence farmers’ enthusiasm and capacity regarding the shift to organic agriculture. Phase Two aimed to examine what has been achieved concerning these concerns and indicate what needs to be done.

### Method of Analysis

The data collected from farmers’ responses were analyzed using qualitative data. The replies were each analyzed individually to capture the frequencies and challenges most often. Answers provided by respondents were grouped according to major concerns, Technical and Cultural/Social, Marketing and Economic, Institutional/Regulatory and Legislative, and Information and Advisory Services. This thematic approach enables an elaboration of views at the current time based on what was generated from phase one of the study. **Table 10** shows the farmer responses is still evident today; twenty years later, organic farming is still challenging. Concerns that need to be addressed include ambiguity of practices, absence of demonstration sites, expensive inputs, limited government support, and weak extension education. Practical training and instruction, market assurance, and subsidies that farmers demand demonstrate that the system is not ready for organic farming, which hinders farmers from adopting this process.

**Table 11** indicates farmer report that the technical, cultural, marketing, and advisory constraints to organic farming are still evident today. These include issues related to the soil’s ability to support sustainable production, pests and diseases, and other inputs. Cultural barriers entice consumers due to high prices, while economics and marketing issues are still unsolved.

### The observed decline in Jordan’s organic farming area

As mentioned in the introduction, there is currently no research addressing the root causes of the notable drop in the adoption of organic farming that has been seen since 2013. Farmers’ opinions on the causes of the decline were gathered through a poll to investigate this matter.

Every respondent highlighted the obstacles noted in Phases One and Two of this study and ascribed the drop to the lack of a national organic agricultural plan. According to the findings, removing these obstacles and putting in place long-term governmental plans are essential to attaining the adoption of sustainable organic farming.

### Contribution of Phase Two Findings

The results obtained in Phase Two advance the knowledge in tackling the perennial issues of adopting organic farming. Validating the impediments revealed in Phase One, namely technical, economic, and advisory in its structure, this phase underlines the systemic character of the barriers. It demonstrates that policy shifts during the last two decades are insufficient. It also defines changing needs, such as rising costs of organic inputs and lack of functional tolls demonstration sites to make subsequent interventions more relevant. In addition, the research points to a lack of implementation of existing policies by creating an organic department that does not cater to critical areas such as certification or practical assistance. The outcomes of these cases serve as a basis for policy recommendations, noting the importance of subsidies, easy certification, and market guarantees. Also, it contributes to theory development and provides a basis on which the following research and intervention on gender mainstreaming in the country should build, calling for the creation of more tailored training platforms and enhanced market connections. Phase Two presents practice-oriented insights for successfully implementing organic farming and harmonizing policy instruments with farmers’ needs.

## DISCUSSION

From the results of phase one, the knowledge and information about organic farming among Jordanian farmers is extremely low, as we reached only 35% of them. This highlights a significant gap in that much information and best practices are poorly communicated or demonstrated. The primary myth regarding organic farming equaling it to manure can be attributed to language and culture barriers, where organic is limited. Further, it revealed that Farmers’ involvement in the focus group discussions provided the understanding that contributed to such knowledge deficit and called for improved health communication (Crowder et al., 2010). The identified barriers were primarily technical, and all the participants perceived higher pest and disease pressure and sizable yield losses. Concerning other challenges, a high % of respondents, 94%, faced poor soil fertility, and 92% had limited access to biological control agents. The study draws similar conclusions, though local to the region, while stressing drought’s extra risk and impacts on the arid lands’ organic farming climates. Although the interviewees did not directly forbid the approach, perceived risk and general social and cultural factors created hesitation and barriers to failing and experimenting with organic farming. This fear, originating from a desire to retain one’s social status and not be embarrassed like in this same community, affirms its validity in other parts of the world, lest the research only be considered a regional-specific work.

While some of these barriers remained the same as earlier identified, including the issue of market access and the relatively high costs of practicing organic farming, new concerns started to arise due to the expansion of the organic foods market and the growing awareness of the impacts of conventional agriculture on the environment (Reganold & Wachter, 2016). This change is in tandem with the changing societal perspective toward going green and minimizing the use of chemicals in agricultural production. However, as Muller et al. noted, organic farming systems face structural and policy-related barriers such as lack of government support and inadequate resource acquisition (Muller et al., 2017). Additionally, the participants noted that despite the environmental benefits of the shift, it was said that the change must be preceded by an appropriate education and training mechanism necessary to enable farmers to embrace sustainable farming practices. Therefore, efforts to balance economic and ecological factors to create a sustainable approach to organic farming and increase the acceptance rate among conventional farming communities will be crucial (Muller et al., 2017)

According to phase two results, two decades later, such perceptions are very similar across farmers. They have also pointed out indefinite misunderstandings and doubts regarding the sustainability of the organic farming concept and its applicability. Purveyors remained utterly constricted by confusion about the rules, insufficient tangible assistance, and persistently high costs. Interestingly, they argued that subsidies, training workforce training, and government setting up demonstration sites should be what the government should do (Muller et al., 2017). Some aspects have become more institutionalized than others, but new problems have been created, specifically regarding certification and inspection. Another proven hindrance is financial: They lack appropriate premium pricing and, more importantly, market access. The fixed and recurrent technical, economic, and informational constraints argue the need for an all-embracing strategy that combines education, financial incentives, and supportive infrastructure to encourage organic farming in arid zones.

## CONCLUSION

Previous literature has provided invaluable insights into evolving challenges and opportunities in organic farming in the last two decades. Further, it showed the importance of sustained research to amend and implement the existing policies and practices that not only benefit Jordan but also serve as a model for other countries with similar agricultural and environmental conditions.

In conclusion, the study highlights diverse barriers impeding the widespread implementation of organic farming in dry regions of Jordan, encompassing technical, economic, perceptual, institutional, and regulatory challenges. Issues such as pest infestation, yield reduction, unclear perceptions of ‘organic farming,’ fragmented institutions, and regulatory gaps hinder the shift. Despite these obstacles, there is potential in the region due to available land and favourable conditions. These challenges call for collaboration with identified stakeholders to define and address issues and simplify the legal framework to promote the efficiency of organic farming in Jordan for sustainable agriculture, environmental conservation, and socioeconomic development.

## Supporting information

survey questions phase 1

survey questions phase 2 table 10&11

## Appendix

**Table 1:** Jordan’s Organic Land Areas 2022.

| Item | Organic area [ha] | Organic share [%] | Area fully converted [ha] | The area under conversion [ha] |
| --- | --- | --- | --- | --- |
| Total organic agri. land 2022 | 1'478 | 1% | 1'478 | - |
| Total organic agri. land 2021 | 1'446 | 1.6% | 1'446 | - |
| 1-year growth | 32 | - | - | - |
| 10-year growth | -1'420 |  |  |  |
| Cereals | 18 | 0.04 | 18 | 0 |
| Coffee | - | - | - | - |
| Temperate fruit | 144 | 2.3% | 144 | 0 |
| Tropical/Subtropical fruit | 5 | 0.1 | 5 | 0 |
| Grapes | 7 | 0.2 | 7 | 0 |
| Olives | 386 | 0.6 | 386 | 0 |
| Vegetables | 39 | 0.1 | 39 | 0 |
| Agricultural land and crops, no details [ha] | 863 |  | 863 | 0 |

Source: derived from FIBL and IFOAM (2024) and Willer, H (2024)

**Table 2:** Socioeconomic key organic indicators.

| Item | Number | Export to EU [MT] | Export to USA [MT] |
| --- | --- | --- | --- |
| Organic producers | 16 |  |  |
| Export | - | 70 | 16 |
| Processors | 7 |  |  |
| Coffee exported MT | 16 |  |  |

Source: derived from FIBL and IFOAM (2024) and Willer, H (2024)

**Table 3:** Key organic indicators for Jordan vs. the Gulf Cooperation Council (GCC) countries.

| Country | Organic area [ha] | Organic share | Organic producers [no.] | Organic retail sales [Million €] | Export to EU and USA [MT] |
| --- | --- | --- | --- | --- | --- |
| Jordan | 2077 | 1% | 16 | - | 86 |
| Oman | 7 | 0.0005% | 1 |  |  |
| Saudi Arabia | 23,315 | 0.01% | 512 | 325 |  |
| United Arab Emirates | 5,419 | 1.4% | 152 |  | 266 |
| Bahrain |  |  | 1 processor |  | 515 |
| Kuwait | 25 | 0.02% | 1 processor |  | 1 exporter |

Source: derived from FIBL and IFOAM (2024) and Willer, H (2024)

**Table 4:**
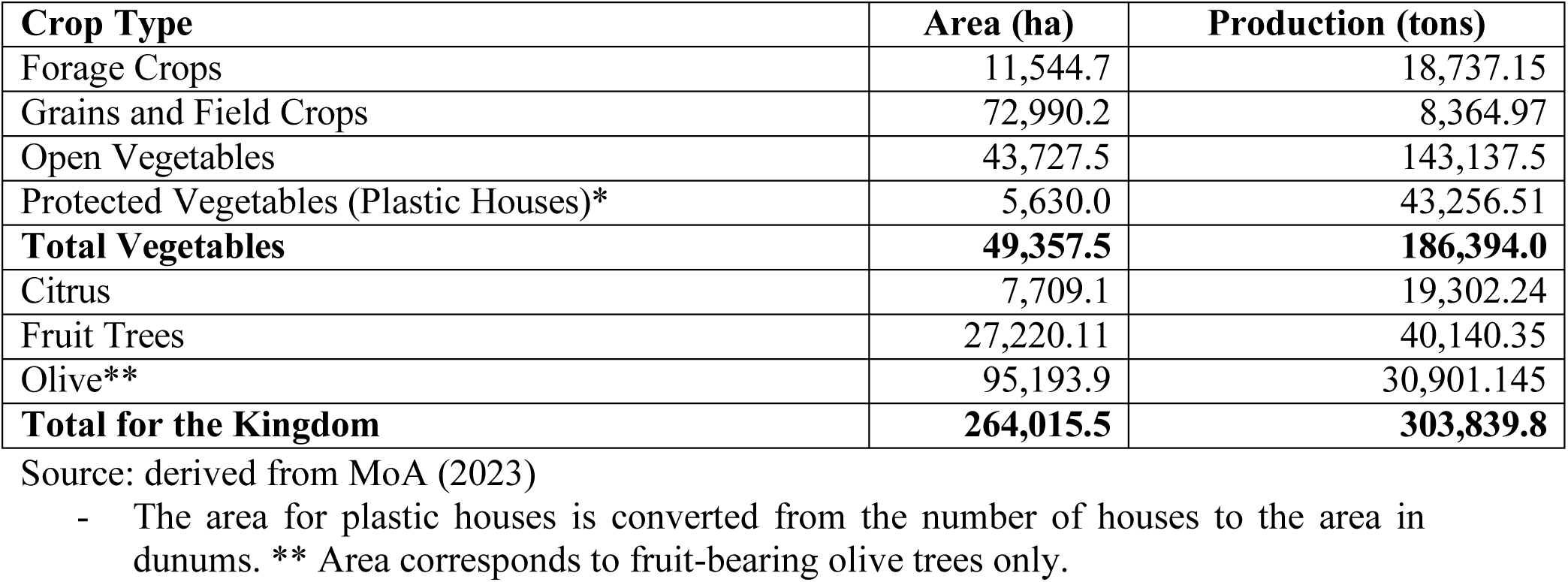
Crop Types, area and production.

**Table 5:** Technical barriers according to the farmers’ response (n=46).

| **Technical barrier** | **Farmers (n=46)**<br>**%** | **Focus groups (n=5)**<br>**%** |
| --- | --- | --- |
| Higher diseases and pest infestation | 100 | 100 |
| Yield reductions (quantity and quality) | 100 | 100 |
| Decrease in plant growth | 91 | 80 |
| Soil fertility is poor | 91 | 60 |
| Risk of trying | 70 | 60 |
| Varieties are not suitable | 61 | 100 |
| Biological control is not available | 61 | 100 |
| Long production periods | 57 | 100 |
| Ignorance of the farmers' experience (knowledge) | 50 | 100 |
| Weather fluctuations | 28 | 40 |

**Table 6:** Economic barriers to adopt of organic farming (n=46).

| **Economic barriers** | **%** | **Frequency** |
| --- | --- | --- |
| There are no market or premium prices for organic farming products | 97.8 | 45.10 |
| Significant investment has already been made | 76.1 | 35.20 |
| Financial commitments | 52.2 | 23.89 |
| Jobs will decrease | 83 | 38.18 |

**Table 7:** Factors favored by organic production in the study area Response of farmers (n=46) and the focus groups (n=5).

| **Response** | **Farmers %** | **Number of focus group %** | **Frequency of Focused Group** |
| --- | --- | --- | --- |
| Wide area and available virgin lands | 93 | 5 | 0.25 |
| Availability of water with good quality | 87 | 5 | 0.25 |
| It is easy to isolate farms in the Badia because the area is vast. | - | 4 | 0.2 |
| Long production season | 87 | 5 | 0.25 |
| Good weather and not polluted | 74 | 4 | 0.2 |
| Soil is not polluted | 83 | 4 | 0.2 |
| Livestock present | 7 | 2 | 0.1 |
| Labor (workers) available | - | 2 | 0.1 |
| The biggest tomato factory in the country | 43 | 2 | 0.1 |
| Farmers accept new profitable ideas | 78 | 3 | 0.15 |

**Table 8:** Types of farms and their cultivation area.

| **Farm data** | **Fruit** | **Mixed** | **Vegetable** | **Total** |
| --- | --- | --- | --- | --- |
| Number of owners | 10 | 12 | 24 | 46 |
| Number of farms | 14 | 14 | 29 | 57 |
| Percentage | 24 | 26 | 50 | 100 |
| Cultivated area ha | 396 | 542 | 551 | 1489 |
| Total land area ha | 425 | 882 | 846 | 2153 |

**Table 9:**
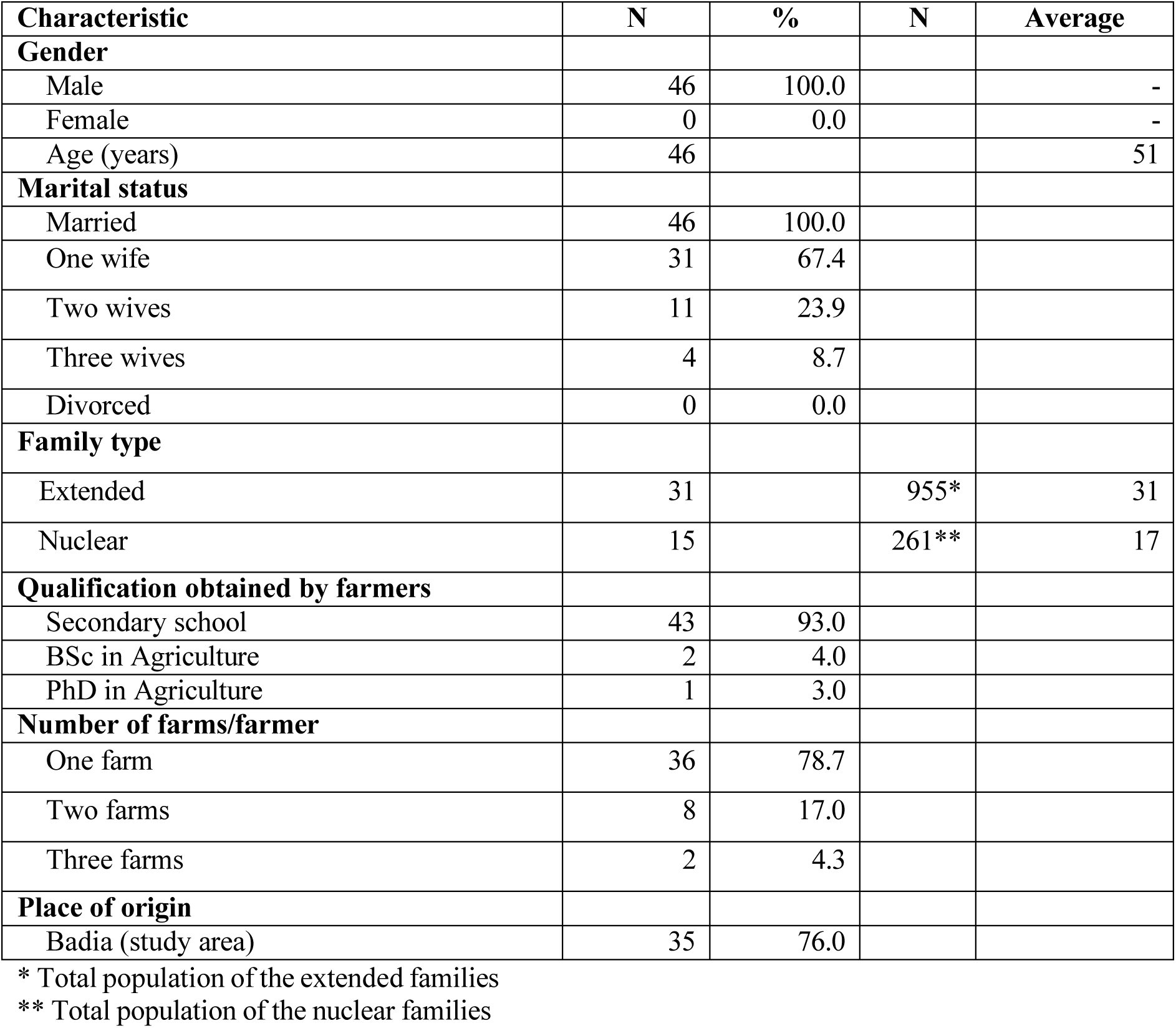
Socio-demographic characteristics of farmers interviewed in the study (n=46).

| **Characteristic** | **N** | **%** | **N** | **Average** |
| --- | --- | --- | --- | --- |
| **Gender** |  |  |  |  |
| Male | 46 | 100.0 |  | - |
| Female | 0 | 0.0 |  | - |
| Age (years) | 46 |  |  | 51 |
| **Marital status** |  |  |  |  |
| Married | 46 | 100.0 |  |  |
| One wife | 31 | 67.4 |  |  |
| Two wives | 11 | 23.9 |  |  |
| Three wives | 4 | 8.7 |  |  |
| Divorced | 0 | 0.0 |  |  |
| **Family type** |  |  |  |  |
| Extended | 31 |  | 955* | 31 |
| Nuclear | 15 |  | 261** | 17 |
| **Qualification obtained by farmers** |  |  |  |  |
| Secondary school | 43 | 93.0 |  |  |
| BSc in Agriculture | 2 | 4.0 |  |  |
| PhD in Agriculture | 1 | 3.0 |  |  |
| **Number of farms/farmer** |  |  |  |  |
| One farm | 36 | 78.7 |  |  |
| Two farms | 8 | 17.0 |  |  |
| Three farms | 2 | 4.3 |  |  |
| **Place of origin** |  |  |  |  |
| Badia (study area) | 35 | 76.0 |  |  |

\* Total population of the extended families
\*\* Total population of the nuclear families

**Table 10:** Farmers’ responses to perception if it still perceived as a barrier after 2 decades of Phase 1 results.

| Respondent | Response |
| --- | --- |
| Farmer 1 | Organic farming is unclear to most farmers and me in my area. I have heard of organic farming, but I am unsure how to implement it sustainably without using chemicals. Still not well explained to farmers. Still a barrier |
| Farmer 2 | Yes, it is not clear to us, and it is just a trend. I have been farming for years, and I know what works. I will not change my methods unless I guarantee they will work. Even there are no demo sites to show us how it works. Still a barrier |
| Farmer 3 | For me, Yes, as a vegetable farmer, but more work must be done to clarify this for all farmers. I am interested in organic farming but worried about the costs. Organic inputs are expensive, and I do not know if they will work. Still a barrier |
| Farmer 4 | No, it is not. The government gives some information, but it is not practical. I use traditional practices, but these cannot be accepted unless my whole farm is certified organic. Still a barrier |
| Farmer 5 | very little and still needs more extension work, especially regarding inspection and certification. It is still a barrier. |
| Farmer 6 | still not clear, and I need more information and support to get started. I am open to organic farming if there is a good market for organic products. However, I need to be sure I can get a fair price for my crops." Still a barrier |
| Farmer 7 | It is not clear at all. The government needs to do more to support organic farming. We need subsidies, training programs, and access to organic inputs. Still a barrier |

**Table 11:**
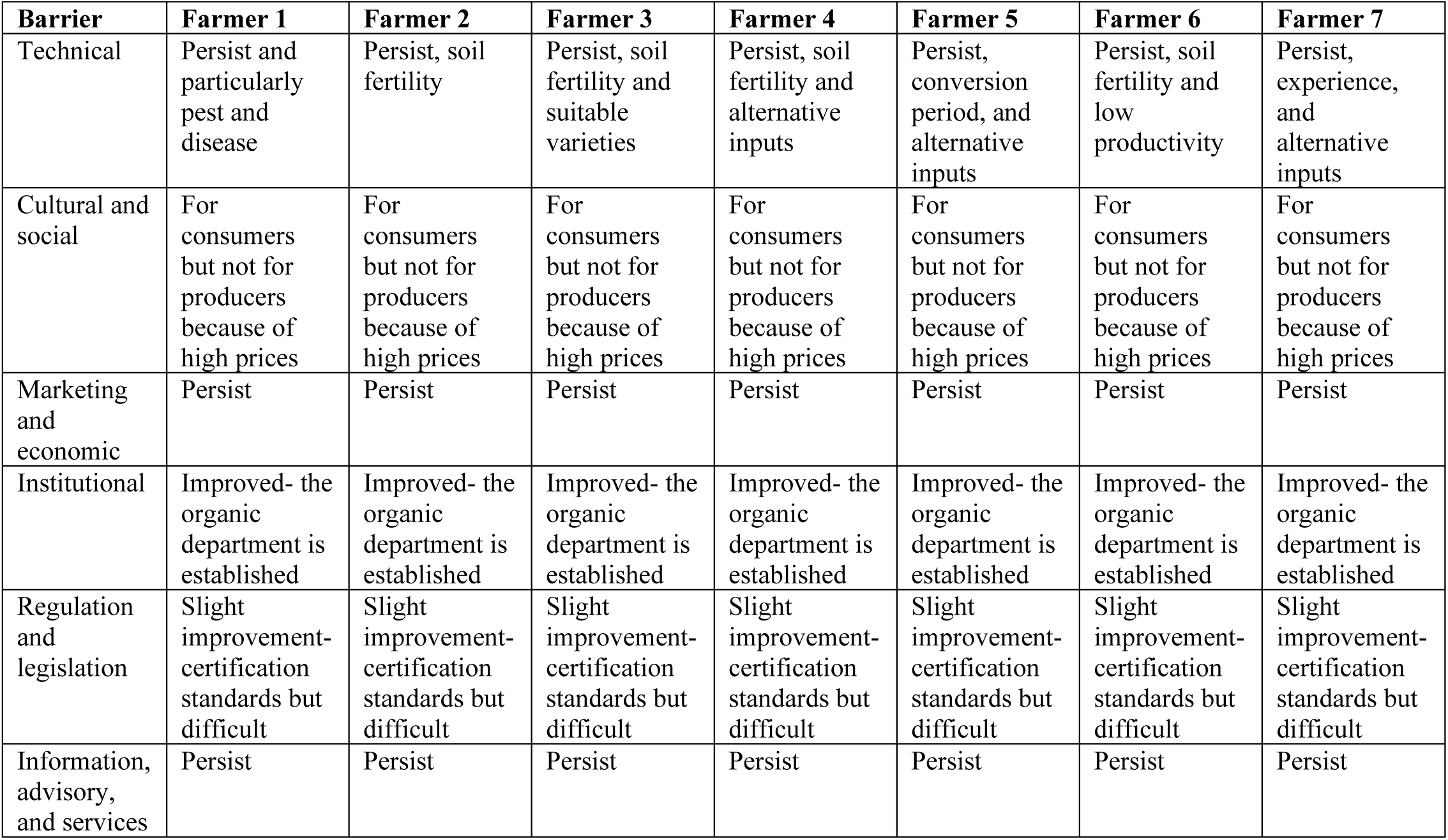
Farmers’ responses to other barriers if they still perceived as barriers after 2 decades of Phase 1 results.

## Notes

### Competing Interest Statement

The authors have declared no competing interest.

