## Supplementary material for "A 2-decade Study of Barriers to the Adoption of Organic Farming in Arid Lands of Jordan": survey questions phase 1

### Supplementary Material– Phase One Questionnaire (Anonymized)

This supplementary file provides the survey instruments and interview guides used in Phase One of the study. All personal identifiers (names, numbers, dates, times, GPS coordinates, and location-specific identifiers) have been removed to ensure confidentiality and compliance with bioRxiv public dissemination requirements.

#### Appendix B. Farmers' Questionnaire (Questions Only)

Sex:
Age:
Family size:
Educational qualification:

A. Farm Information
- Total farm area (ha)
- Crops grown
- Livestock (type and number)
- Jobs existing on the farm and wages paid
- Family members depending on on-farm income
- Family members depending on off-farm income

B. Extension
- Best way to obtain agricultural information

C. Pest Control and Management
- Main pests affecting crops and control methods used
- Types of pesticides used (ranked by relative frequency)
- Non-chemical pest control strategies (Checklist A)

D. Soil Fertility Management
- Land preparation steps (soil analysis, moisture, pH, amendments)
- Inorganic fertilisation programme from land preparation to harvest

E. Organic Fertilisers
- Types used (type, amount per unit area, price)
- Reasons for use
- Constraints to use
- Non-chemical soil fertility strategies (Checklist B)

F. Environmental Impacts
- Environmental impacts observed from conventional farming practices

G. Organic Farming
- Awareness and understanding of organic farming
- Technical barriers
- Cultural barriers
- Economic barriers
- Potential enabling factors

H. Adoption of Organic Farming
- Willingness to adopt organic farming if advised by authorities and reasons

#### Checklist A. Non-Chemical Pest Control Strategies

Sanitation and hygiene; crop rotation; fallow periods; hand picking; live barriers; mulching (black plastic); resistant varieties; crop patterning; water spraying; summer and winter oils; tillage and irrigation management; timely planting; sulphur; weeding; lime (white stone).

#### Checklist B. Non-Chemical Soil Fertility Strategies

Compost application; manure; crop rotation; fallow periods; mineral rocks; mulching; timing of agricultural activities (seeding, fertilisation); windbreaks.

#### Farm Observation Checklist

Water source; water reservoir; water pumping system; irrigation system; fertilising systems; land preparation; planting; pest control; storage of pesticides and fertilisers; fruit collection; packaging and marketing; farm layout.

#### Private Agricultural Store Suppliers’ Questionnaire (Questions Only)

- Organic pesticides sold (type, pest, crop, source)
- Organic fertilisers sold (type, crop, source)
- Observed technical and cultural barriers to organic farming
- Potential enabling factors for organic production

#### Key Informant Interview Guides (Roles Only)

Participants:
- Senior academic (agriculture)
- Senior agricultural policymaker
- Agricultural input or IPM business representative

Topics: perception of organic farming, institutional barriers, market experience, policy environment, and recommendations.

#### Decision-Maker Interview Guides

Participants:
- Agricultural regulatory unit
- Senior ministry leadership
- Agricultural policy unit

Topics: legislation, standards, certification systems, farmer involvement, and future planning for organic agriculture.
