## Supplementary material for "A 2-decade Study of Barriers to the Adoption of Organic Farming in Arid Lands of Jordan": survey questions phase 2 table 10&11

### Supplementary Material– Phase 2 Farmer Interview Questions

This supplementary document presents the Phase 2 interview questions used to explore whether barriers to organic farming identified during Phase 1 (approximately two decades earlier) are still perceived by farmers. The questions were asked through semi-structured interviews. All questions were originally conducted in Arabic and later translated into English. No personal identifiers were collected.

#### Section 1. Farmers' Perception of Organic Farming

Q1. How clear and understandable is the concept of organic farming for you and other farmers in your area? Please explain.

Q2. Do you think organic farming can realistically be applied on your farm under current conditions? Why or why not?

Q3. Compared with about 20 years ago, do you feel that farmers' understanding of organic farming has improved, stayed the same, or declined? Please explain.

Q4. Under what conditions would you personally consider adopting organic farming practices?

Q5. Overall, do you still consider farmers' perception and understanding of organic farming to be a barrier to adoption today?

#### Section 2. Technical Barriers

Q6. Do technical challenges such as pest and disease management, soil fertility, crop productivity, availability of suitable varieties, or access to alternative inputs still limit organic farming adoption? Please specify.

Q7. Compared with the past, have these technical challenges improved, remained the same, or worsened?

#### Section 3. Cultural and Social Factors

Q8. Do cultural or social factors discourage farmers from producing organic crops? Please explain.

Q9. Do consumers' attitudes or willingness to pay higher prices influence farmers' decisions to adopt organic farming?

#### Section 4. Marketing and Economic Barriers

Q10. Is there, in your opinion, a reliable and accessible market for organic products at present?

Q11. Do production costs, price uncertainty, or financial risks discourage farmers from adopting organic farming?

#### Section 5. Institutional Support

Q12. Do you believe government institutions currently provide better support for organic farming than in the past? Please explain.

Q13. Have institutional structures related to organic farming improved over the last two decades?

#### Section 6. Regulation and Legislation

Q14. Are current organic certification standards and regulations supportive or restrictive for farmers? Please explain.

Q15. Compared with the past, do you think regulations related to organic farming have improved?

#### Section 7. Information, Advisory, and Extension Services

Q16. Do farmers currently have sufficient access to information, training, and advisory services related to organic farming?

Q17. In your opinion, do extension officers have adequate knowledge and capacity to support organic farming adoption?
